# Gastruloids as *in vitro* models of embryonic blood development with spatial and temporal resolution

**DOI:** 10.1101/2021.03.21.436320

**Authors:** Giuliana Rossi, Sonja Giger, Tania Hübscher, Matthias P. Lutolf

**Affiliations:** Laboratory of Stem Cell Bioengineering, Institute of Bioengineering, School of Life Sciences and School of Engineering, École Polytechnique Fédérale de Lausanne (EPFL), Lausanne, 1015, Vaud, Switzerland; Institute of Chemical Sciences and Engineering, School of Basic Science, École Polytechnique Fédérale de Lausanne (EPFL), Lausanne, 1015, Vaud, Switzerland

## Abstract

Gastruloids are three-dimensional embryonic organoids that reproduce key features of early mammalian development *in vitro* with unique scalability, accessibility, and spatiotemporal similarity to real embryos. Recently, we adapted gastruloid culture conditions to promote cardiovascular development. In this work, we extended these conditions to capture features of embryonic blood development through a combination of immunophenotyping, detailed transcriptomics analysis, and identification of blood stem/progenitor cell potency. We uncovered the emergence of blood progenitor and erythroid-like cell populations in late gastruloids and showed the multipotent clonogenic capacity of these cells, both *in vitro* and after transplantation into irradiated mice. We also identified the spatial localization near a vessel-like plexus in the anterior of gastruloids with similarities to the emergence of blood stem cells in the embryo. These results highlight the potential and applicability of gastruloids to the *in vitro* study of complex processes in embryonic blood development with spatiotemporal fidelity.

## Introduction

Mammalian embryos develop in the uterus and are dependent on maternal interactions, which raises scientific and ethical challenges in accessing them for developmental studies. Embryonic organoids are 3D models that are experimental alternatives to mammalian embryos and offer the unprecedented potential to study aspects of embryogenesis *in vitro*. Due to their accessibility, scalability, and experimental versatility, embryonic organoids offer promising alternatives and complements to studies in animal models (Harrison et al., 2017; Rivron et al., 2018; Rossi et al., 2018; Shao et al., 2017a, 2017b; Sozen et al., 2018; van den Brink et al., 2014; Zheng et al., 2019). Gastruloids, a type of embryonic organoid, are aggregates of embryonic stem cells (ESCs) that mimic aspects of post-implantation development, such as symmetry breaking, gastrulation and establishment of the three major body axes, when cultured under the correct conditions (Beccari et al., 2018; van den Brink et al., 2014).

We have recently shown that gastruloid culture conditions can be steered to promote early cardiovascular development, or the formation of what resembles a vascular network, and a cardiac primordium (Rossi et al., 2020). Cardiovascular development is connected with blood emergence and early blood development depends on the endothelial-to-hematopoietic transition (ETH), a process in which vascular cells of the hemogenic endothelium progressively lose their endothelial signature and activate a hematopoietic transcriptional program (Jaffredo et al., 1998; Ottersbach, 2019; Zovein et al., 2008). Hematopoietic system development occurs in two successive, spatially and temporally restricted waves (Costa et al., 2012). Primitive hematopoiesis begins around embryonic day 7.5 (E7.5) in the yolk sac blood islands, which is defined by the initial wave of blood cell production before circulation is established (Maximow, 1924; Moore and Metcalf, 1970). After establishment of circulation, definitive hematopoiesis takes place from E8.5 to E10.5 at various embryonic sites: the placenta, the aorta-gonad-mesonephros (AGM) region, and the umbilical and vitelline arteries (de Bruijn et al., 2000; Gekas et al., 2005; Medvinsky and Dzierzak, 1996; Müller et al., 1994). During later embryonic development, the fetal liver is colonized through circulation, and becomes the main organ for hematopoietic stem cell (HSC) expansion and maturation (Ema and Nakauchi, 2000; Morrison et al., 1995). Around E16.5, hematopoietic progenitors from the fetal liver start to colonize the developing bones to form the classical adult hematopoietic stem cell niche (Chotinantakul and Leeanansaksiri, 2012; Christensen et al., 2004). Due to the endothelial origin of blood cells, we hypothesized that we could capture the early stages of blood development in gastruloids.

Here we show that gastruloids cultured in cardiovascular-inducing conditions (Rossi et al., 2020) display a hematopoiesis-related transcriptional signature and express surface markers characteristic of early hematopoietic cells. Specifically, we describe the emergence of a population of cells with CD34, c-Kit, and CD41 markers, which correspond to an early blood progenitor phenotype. We also describe a population of Ter-119^+^ cells that display an erythroid phenotype. Analogously to embryonic development, we observed that blood progenitors accumulate from 96 h to 168 h in gastruloids and, at 168 h, we observed their multilineage clonogenic potential both *in vitro* and *in vivo*. We also discovered that these cells are located anteriorly, in proximity to the vascular-like plexus, in line with observations in mammalian embryos. Taken together, these data suggest that gastruloids can model the earlier stages of hematopoietic development and have great utility as an *in vitro* model system to study mechanisms of blood development.

## Results

### Gastruloids display a hematopoiesis-related transcriptional signature

We recently described a protocol to promote cardiovascular development in gastruloids, which required the addition of VEGF, bFGF, and ascorbic acid to standard gastruloid growth culture conditions (Rossi et al., 2020). As VEGF and bFGF are necessary for hematopoiesis *in vitro* and *in vivo* (Daniel et al., 2016; Irion et al., 2010; Kennedy et al., 2007; Mikkola et al., 2003; Motazedian et al., 2020; Pardanaud and Dieterlen-Lièvre, 1999; Sugimura et al., 2017), we hypothesized that these culture conditions could also support the development of the blood lineage. Thus, we mined the scRNA-seq dataset of gastruloids grown in the presence of VEGF and bFGF (Rossi et al., 2020), with a focus on clusters that were on the trajectory from epiblast to endothelial progenitors (**Figure 1A**). From gene expression analysis of cells in these clusters, we noted the progressive emergence of a population that expressed characteristic markers of blood progenitors. Around 96 h (**Figure 1B**), we observed a population of cells expressing *Brachyury*, *Mixl1*, *Pdgfra,* and *Kdr/Flk1/Vegfr2*, which was indicative of mesoderm patterned to a hematopoietic fate (Ivanovs et al., 2017) (**Figure 1C**). Co-expression of *Kdr* and *Brachyury* is known to mark a common progenitor for cardiac, endothelial, and hematopoietic lineages (Huber et al., 2004; Kattman et al., 2006). We also noted that *Sox17*, which marks the hemogenic endothelium *in vivo* (Clarke et al., 2013), was upregulated over time, along with transcription factors and surface antigens typically associated with hematopoietic development, such as *Tal1/SCL*, *Etv2*, *Lmo2*, *Thy1*, and *Gata2* (**Figure 1C**) (Gao et al., 2013; Koyano-Nakagawa et al., 2012; Pater et al., 2013; Porcher et al., 1996; Tsai et al., 1994). In later stages of development, around 144 h to 168 h (**Figure 1B**), we observed the start of the expressions of *Kit* and its ligand *Kitl*, *Cd34*, *Itga2b/Cd41*, *Cd93*, *Cdh5/VE-cadherin*, and *Mecom/Evi-1*, which are typical markers of developing blood progenitors (Baumann et al., 2004; Bertrand et al., 2005; Ferkowicz et al., 2003; Goyama et al., 2008; Matsubara et al., 2005; Mikkola et al., 2003; Yoder et al., 1997; Zovein et al., 2008) (**Figure 1C**). In line with previously reported observations (Beccari et al., 2018), the gastruloids also expressed *Hoxb4* (**Figure 1C**), which has been shown to promote hematopoietic differentiation *in vitro* (Kyba et al., 2002). Finally, at 168 h, we observed a small population that expressed the hemoglobin embryonic isoforms *Hbb-y* and *Hbb-bh1* (**Figure 1C**) (Kingsley et al., 2006). Intriguingly, the hemoglobin genes were associated with an endothelial sub-cluster that expressed typical endocardial genes (Rossi et al., 2020), which is suggestive of the emergence from hemogenic endocardium (Nakano et al., 2013). We performed RT-qPCR analysis of gastruloids at different time points, which confirmed the progressive upregulation of hematopoiesis-related factors (**Figure S1A-E**), as well as the expression of *Hoxb4* (**Figure S1F**). We also observed the expression of embryonic (*Hbb-y*, *Hbb-bh, Hba-x*) but not adult (*Hbb-b1*, *Hbb-b2*, *Hba-a1*) hemoglobin isoforms (**Figure S1G-L**). Taken together, these results suggest that gastruloids express key hematopoiesis-related genes in a temporal sequence that follows embryonic development.

**Figure 1.**
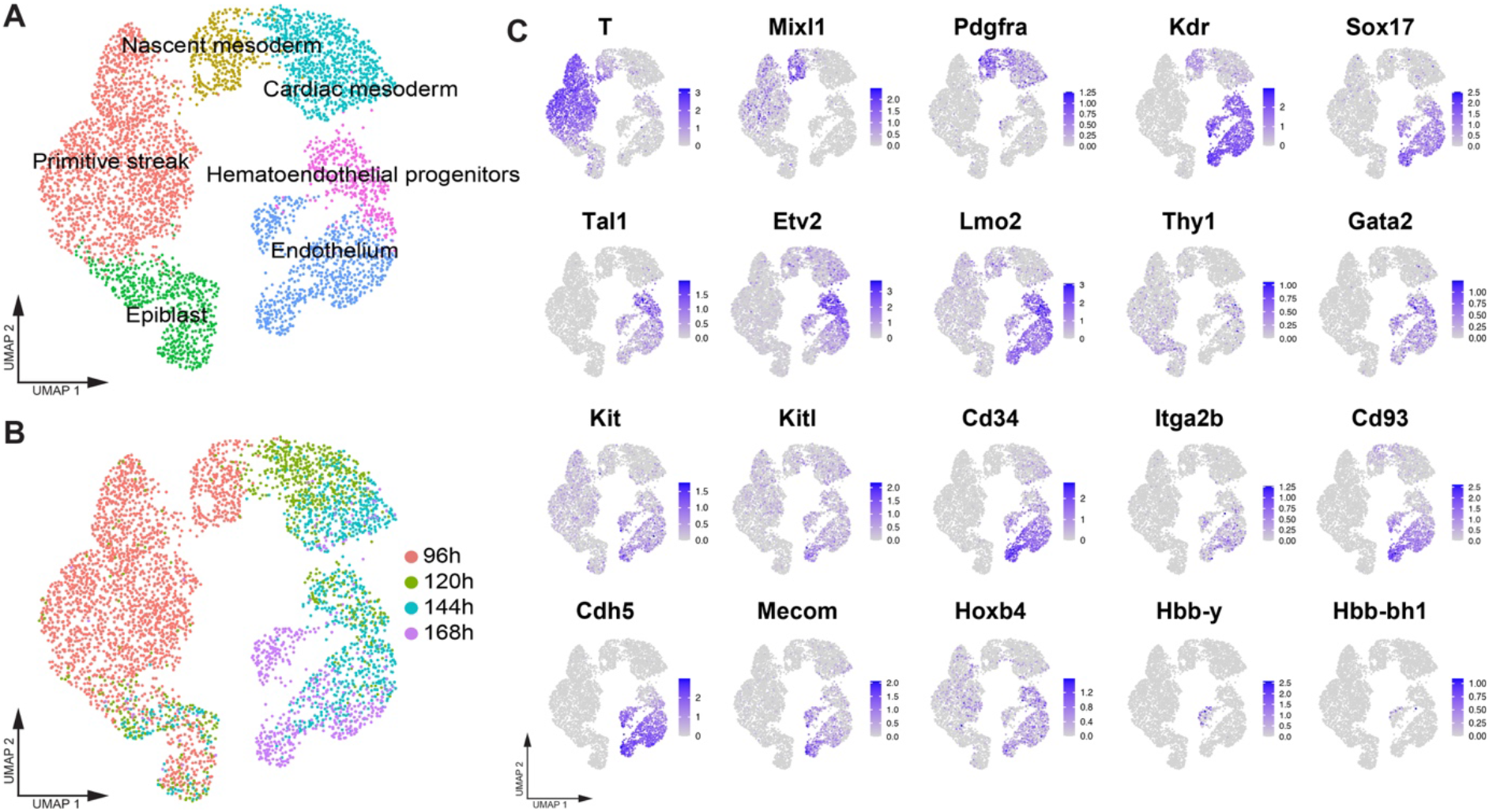
Gastruloids express markers of early blood development. UMAP plots showing (**A**) clusters on the trajectories from epiblast to endothelium and (**B**) the distribution of cells in the UMAP plots according to analyzed time points (**C**) UMAP plots showing the expression of hematopoiesis-related genes in the scRNA-seq dataset. Dataset from (Rossi et al., 2020). n = 2 replicates per time point.

### Early blood cells in gastruloids are identified by surface marker analysis

We can examine the expression of typical surface markers to identify blood progenitors. Towards this goal, we performed FACS analysis of gastruloids from 96 h to 168 h to determine the expression of canonical early hematopoietic markers across three different cell lines: *Sox1-GFP::Brachyury-mCherry* (Deluz et al., 2016), *Flk1-GFP* (Jakobsson et al., 2010), and *Gata6-Venus (Freyer et al., 2015)*. We first observed the expression of the hematopoietic markers CD34, c-kit, and CD93. As a marker of early expression during embryonic development, CD34 was upregulated from 120 h (**Figure S2A**). CD34 also marks vascular and hematopoietic progenitors within the hemogenic endothelium (Wood et al., 1997). c-kit was expressed with some fluctuations throughout the gastruloid culture (**Figure S2B**). While both c-kit and CD34 are consistently expressed in early embryonic hematopoietic cells as well as adult HSCs, their expression is not restricted to these cell types (Baumann et al., 2004; Matsubara et al., 2005; Sugiyama et al., 2011; Wood et al., 1997; Yoder et al., 1997). We noted that CD93 was upregulated before 120 h, and its levels increased progressively over time, most evidently in the *Sox1-GFP::Brachyury-mCherry* and *Gata6-Venus* cell lines (**Figure S2C**). CD93 is expressed early on during embryonic development, in both endothelial cells and hematopoietic progenitors (Petrenko et al., 1999).

Between 144 h and 168 h, we then observed an accumulation of CD41^+^ cells, which is a key marker for the onset of hematopoiesis in the embryo (Ferkowicz et al., 2003; Mikkola et al., 2003); this accumulation was most prominent in the *Gata6-Venus* cell line (**Figure S2D**). Of note, it was previously shown that early hematopoietic progenitors lack the expression of the pan-hematopoietic markers CD45 and Sca1, which are upregulated upon later stages of blood ontogeny (Chotinantakul and Leeanansaksiri, 2012). In line with these *in vivo* findings, CD45^+^ and Sca1^+^ cells emerged only during later stages of gastruloid development (**Figure S2E, F**). Additionally, we looked for the expression of Ter119, an erythroid lineage marker that is upregulated in the embryo from E8.0 in evolving erythroid progenitor cells, with concomitant downregulation of the primitive hematopoietic marker CD41 (Ferkowicz et al., 2003). Ter119^+^ cells emerged in gastruloids around 120 h and were maintained on a similar level until 168 h (**Figure S2G**). Finally, the endothelial markers CD31 and Flk1 were also upregulated starting around 120 h (**Figure S2H, I)**, which correlates with previous reports (Rossi et al., 2020).

The concomitant upregulation of hematopoietic and vascular surface markers in gastruloids suggests the emergence of hematoendothelial progenitors. Although such markers are not unique to hematopoietic cells, they can be defined by combinations of markers. A published assessment of the colony formation potential during early embryonic stages (from E7.0 to E9.5) showed that CD34^+^/cKit^+^/CD41^+^ cells from the yolk sac, AGM, and fetal liver held most of the multipotent capacity (Ferkowicz et al., 2003). Interestingly, we found a consistent upregulation of the cKit^+^/CD34^+^ population in gastruloids from 120 h (**Figure 2A, B**), whereas the cKit^+^/CD34^+^/TER119^−^/CD41^+^ population only appeared after 144 h (**Figure 2C, D**).

**Figure 2.**
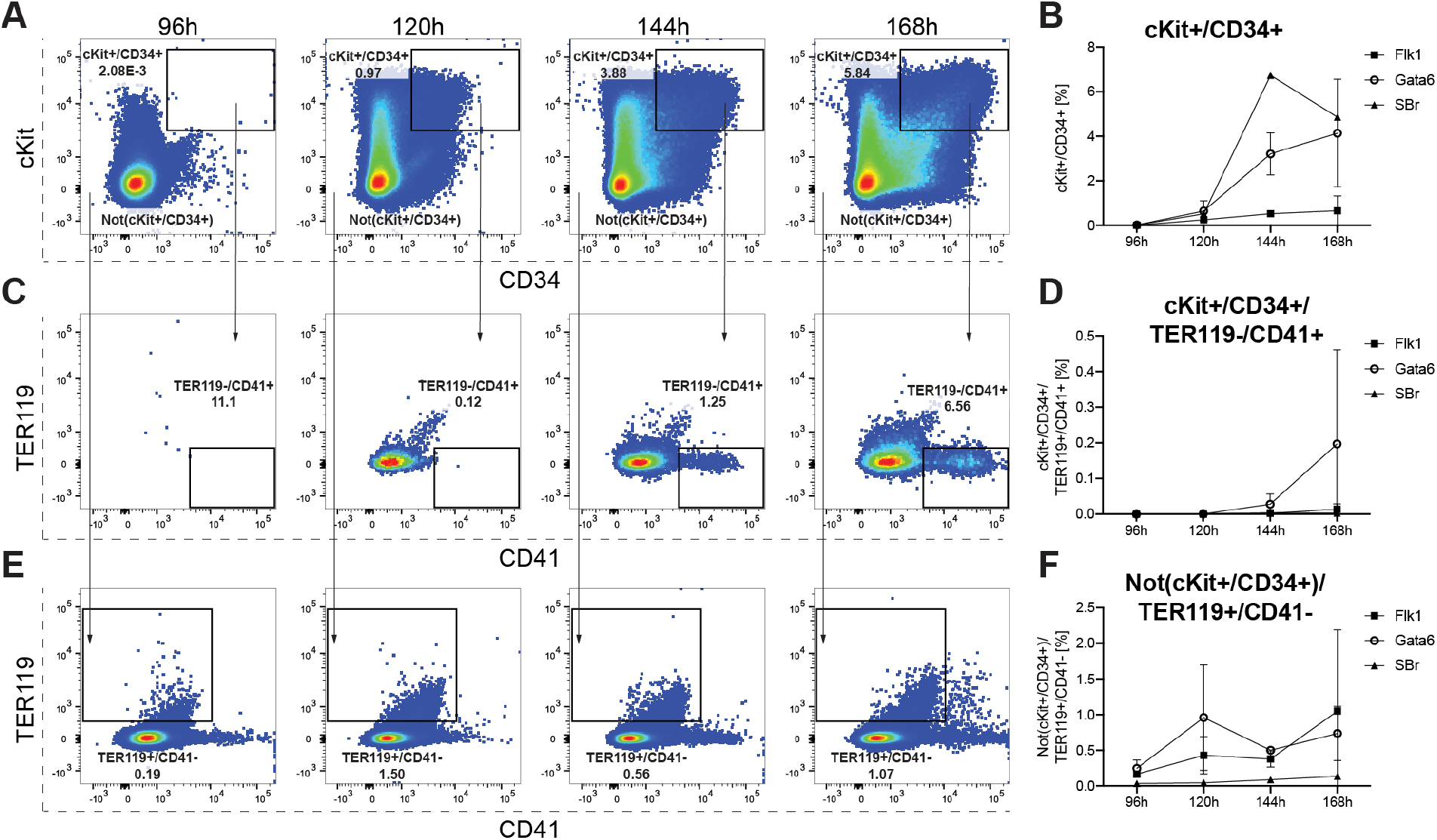
Emergence of hematopoietic and erythroid progenitors in gastruloids. (**A**) Representative FACS plots and (**B**) relative quantification of the emergence of the cKit^+^/CD34^+^ populations in gastruloids from 96 h to 168 h. The cKit^+^/CD34^+^ population was gated further for TER119^−^/CD41^+^ populations, with (**C**) representative FACS plots and (**D**) relative quantification. The Not(cKit^+^/CD34^+^) population was additionally gated for TER119^+^/CD41^−^, with (**E**) representative FACs plots and (**F**) relative quantification. The graphs in **B**, **D**, **F** show the changes in the percentage of positive cells over time. *Flk1-GFP* (n = 2 replicates), *Gata6-Venus* (n = 2 replicates) and *Sox1-GFP::Brachyury-mCherry* (n = 1 replicate). Data are represented as mean ± SD.

The emergence of erythroid cells in the embryo occurs from E8.0 onwards and is marked by the upregulation of the erythroid lineage marker Ter119 with subsequent downregulation of CD41 (Ferkowicz et al., 2003). We observed that Not(cKit+/CD34+)/ Ter119+/CD41-population is upregulated early on and fluctuated over the time course of the gastruloid culture (**Figure 2 E, F**). Our results suggest that primitive hematopoietic precursors and erythroid progenitor cells emerge during late stages of gastruloid development.

### CD41^+^ cells in gastruloids localize near a vascular-like network

Gastruloids display a highly coordinated temporal recapitulation of *in vivo* development and, most importantly, mimic the embryonic organization of tissues in space (Beccari et al., 2018; Rossi et al., 2020). During development, CD41^+^ hematopoietic precursors are found near the Flk1^+^ endothelial network in the most proximal portion of the yolk sac (Ferkowicz et al., 2003). Subsequently, the hemogenic endothelium is found in the AGM region located in the anterior portion of the embryo (Medvinsky et al., 2011). Within the hemogenic endothelium, Sox17 has been shown to play a functional role in HSC development and is a marker for hemogenic endothelial cells and emerging hematopoietic cells (Clarke et al., 2013; Jung et al., 2019). We discovered that Sox17 is expressed in the nuclei of CD31^+^ and CD34^+^ cells, which mark the vascular-like network, and in the anterior region of gastruloids at 168 h (**Figure 3A-D** and **Figure S3A, B**). Moreover, we observed the expression of CD41 in clusters of cells located near the endothelial network that expressed CD31 (**Figure 3E, F** and **Figure S3C**). Taken together, these data suggest that gastruloids support the formation of hemogenic endothelial cells, from which CD41^+^ hematopoietic precursors emerge.

**Figure 3.**
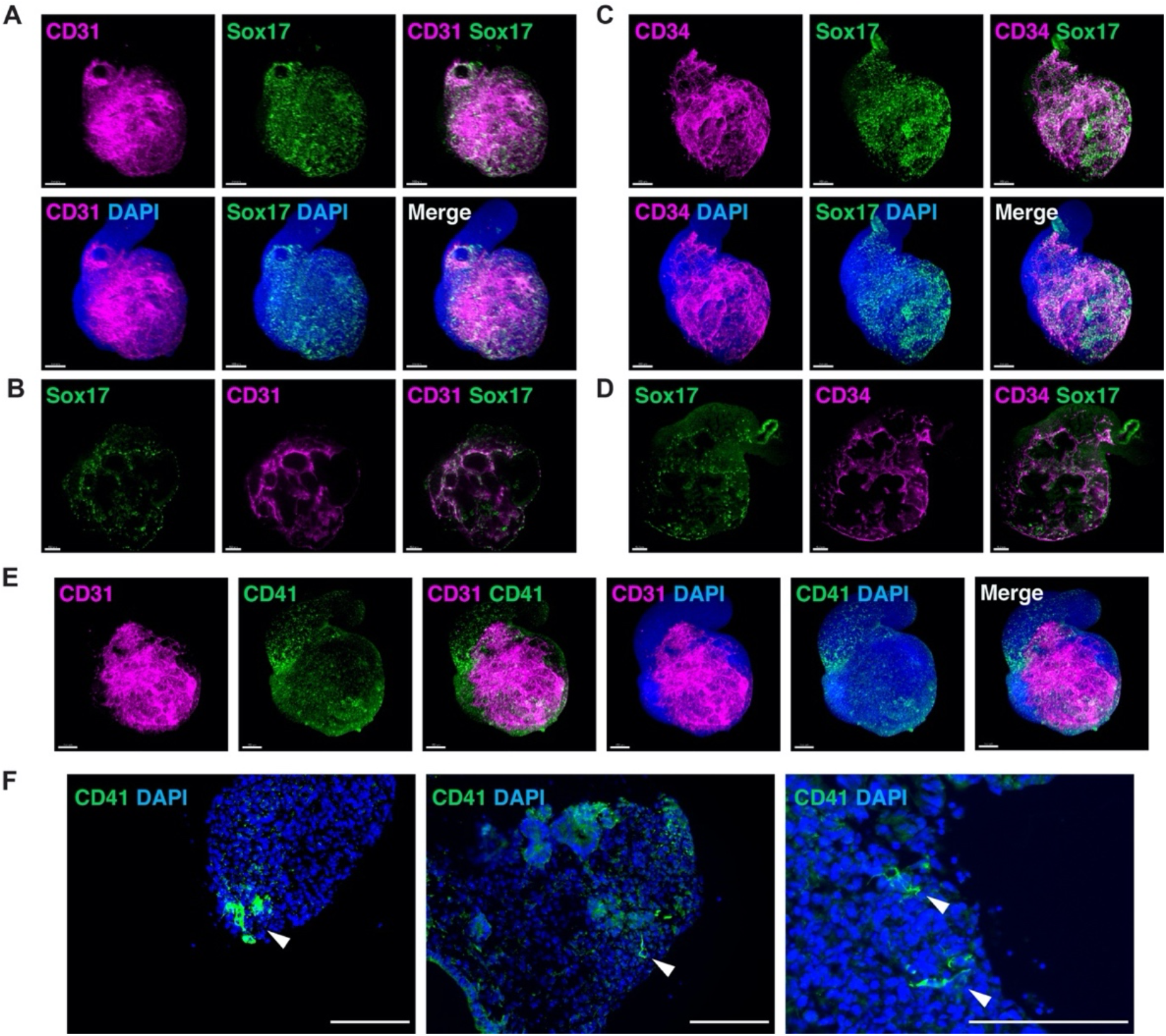
Emergence of blood progenitors in gastruloids. (**A**) Representative light-sheet images of *Gata6-Venus* gastruloids at 168 h showing the co-expression of CD31 and Sox17 in a 3D reconstruction and (**B**) in a single Z-plane. (**C**) Representative light-sheet images of *Gata6-Venus* gastruloids at 168 h showing the co-expression of CD34 and Sox17 in a 3D reconstruction and (**D**) a single Z-plane. (**E**) Representative light-sheet images of *Gata6-Venus* gastruloids at 168 h showing CD41^+^ clusters that arise near the CD31^+^ vascular network. (**F)** Representative sections of *Gata6-Venus* gastruloids at 168 h show CD41^+^ cells at higher magnification. Scale bars,100µm.

### Blood progenitors display multilineage clonogenic potential *in vitro* and *in vivo*

To assess the multipotency of the blood progenitors that arise in gastruloids around 168 h, we isolated two potential populations by FACS: a hematopoietic progenitor population marked by cKit^+^/CD34^+^/TER119-/CD41^+^ and a primitive erythroid population marked by Not(cKit^+^/CD34^+^)/TER119^+^/CD41- (**Figure 4A**). We tested both populations in a colony– forming-unit (CFU) assay. Notably, we observed the formation of morphologically different colonies derived from the cKit^+^/CD34^+^/TER119-/CD41^+^ hematopoietic progenitor population (**Figure 4B**). Colonies were identifiable after 7 days from cell seeding and expanded from day 7 to day 14 (**Figure S4A)**, which suggested that the hematopoietic clones could form efficiently proliferating colonies. Specifically, we observed the formation of colonies that contained single, round-to-slightly elongated cells, which we classified as CFU-megakaryocyte (CFU-Mk); colonies with the typical appearance of CFU-granulocyte, macrophage (CFU-GM), which contained multiple clusters with a dense core of oval-to-round cells with a grey appearance (macrophages) or small, bright and round cells (granulocytes); potential CFU-granulocyte (CFU-G) colonies; and also multipotential-like colonies of CFU-granulocyte, erythroid, macrophage, megakaryocyte (CFU-GEMM), which contained multiple cell types from different lineages (**Figure 4B**). Moreover, we noted the appearance of red-to-brown colonies that indicated the presence of erythroid progenitors, which contained individual erythroid clusters with tiny, irregularly shaped, and often fused cells typical of burst-forming-unit-erythroid (BFU-E) (**Figure 4B**). The Not(cKit^+^/CD34^+^)/TER119^+^/CD41^-^ mainly gave rise to CFU-erythroid (CFU-E) colonies, which were mostly small and contained clusters of irregularly shaped and fused erythroblasts with a similar red-brown color to BFU-E colonies (**Figure 4C**). These colonies were much smaller and did not show evident expansion from 7 to 14 days (**Figure S4B)**. Controls with cKit^+^/CD34^+^/TER119^-^/CD41^-^ and Not(cKit^+^/CD34^+^)/TER119^-^/CD41^-^ populations did not show any evident CFU potential and undifferentiated mouse embryonic stem cells (mESCs) formed only fibroblastic-like clusters (**Figure S4C**). These data support the idea that the expression of CD41 is essential for the establishment of the multipotent potential of hematopoietic progenitor cells in gastruloids, akin to *in vivo* observations (Bertrand et al., 2005; Ferkowicz et al., 2003).

**Figure 4.**
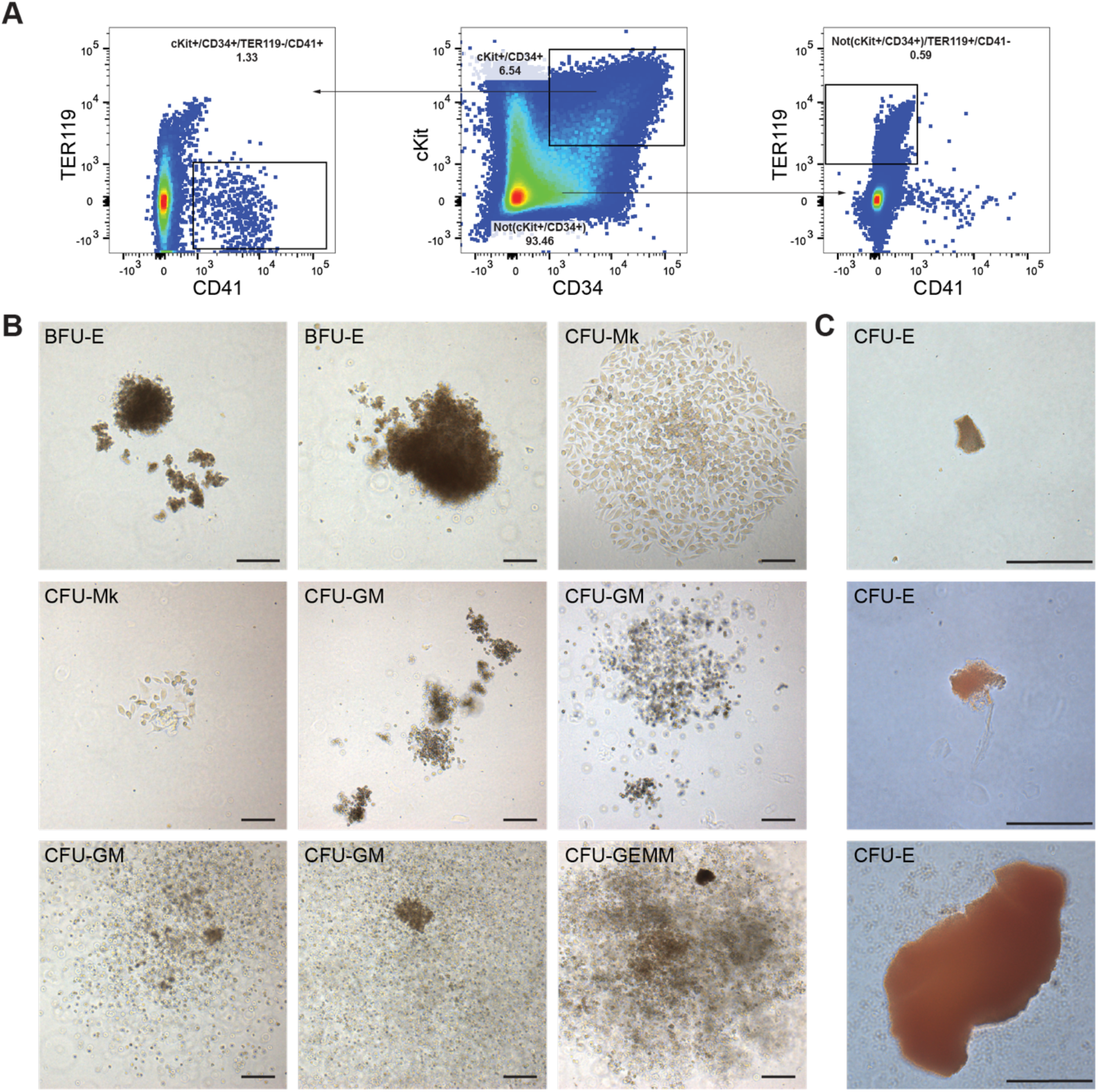
Multilineage clonogenic potential of gastruloid-derived blood progenitors. **(A)** Gating strategy for the isolation of cKit^+^/CD34^+^/TER119^−^/CD41^+^ hematopoietic progenitors and Not(cKit^+^/CD34^+^)/TER119^+^/CD41^−^ primitive erythroid cells from *Gata6-Venus* gastruloids at 168 h. (**B)** Representative images of BFU-E, CFU-Mk, CFU-GM, CFU-E and CFU-GEMM colonies derived from cKit^+^/CD34^+^/TER119^−^/CD41^+^ cells after 17 days of culture in Methocult. (**C)** Representative images of CFU-E colonies derived from the Not(cKit^+^/CD34^+^)/TER119^+^/CD41^−^ population after 17 days of culture in Methocult. BFU-E, burst-forming-unit-erythroid; CFU-Mk, colony forming unit-megakaryocyte; CFU-GM, colony forming unit-granulocyte, macrophage; CFU-GEMM, colony forming unit-granulocyte, erythroid, macrophage, megakaryocyte; CFU-E, colony forming unit-erythroid. Scale bars 200 μm.

Transplantation studies have shown that long-term repopulating HSCs (LTR-HSCs) appear after E10 to E11 in the AGM region of the embryo, a time point that is not attained with current gastruloid culture protocols. However, definitive colony-forming-unit-spleen (CFU-S) can been observed from E9.5 (Medvinsky et al., 2011; Medvinsky and Dzierzak, 1996; Müller et al., 1994). We hypothesized that hematopoietic progenitors within gastruloids at 168 h harbored the potential to generate CFU-S upon transplantation in irradiated animals. As shown in **Figure 5**, we dissociated the gastruloids into single cells at 168 h and injected 5×10^6^ gastruloid cells into the tail vein of lethally irradiated mouse recipients together with 2×10^4^ competitor bone marrow cells. After 11 days, we assayed their capacity to form CFU-S relative to control mice, which were injected with only the competitor bone marrow cells (**Figure 5A**). The presence of CFU-S was significantly higher in the spleens of recipient mice injected with gastruloid cells compared to control mice (**Figure 5B, C**). Altogether, our data indicate that a multipotent hematopoietic progenitor population is developing in gastruloids at 168 h. Surface marker characteristics and the ability to form CFU-S upon *in vivo* transplantation suggest that this population is reminiscent of hematopoietic progenitors that arise on E8.5 to E9.5 in embryonic development.

**Figure 5.**
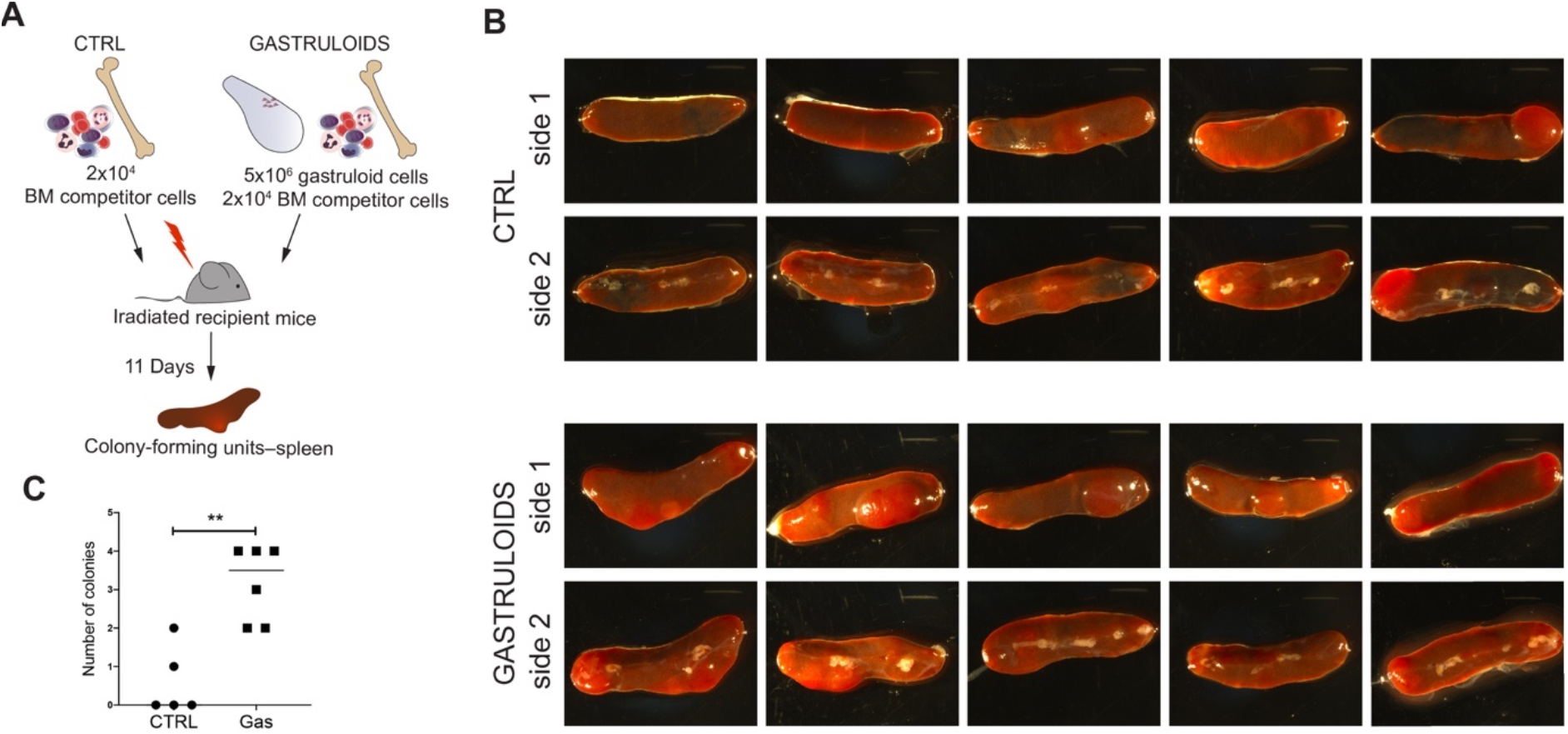
Gastruloid-derived cells form CFU-S upon transplantation. **(A)** Schematic overview of the transplantation strategy and analysis of CFU-S. (**B)** Stereomicroscope images of spleens from 5 different recipient mice imaged on both sides. (**C)** Quantification of the number of colonies observed in recipient mice transplanted with control cells or gastruloids. Single data points are shown in graph. n = 5 control mice and n = 6 mice were injected with gastruloid cells. CTRL, control. BM, bone marrow.

## Discussion

The challenge of studying the development of the hematopoietic system highlights the need for relevant *in vitro* models that can recapitulate the *in vivo* processes with spatial and temporal fidelity. We have recently shown that progenitor cells generated during gastruloid development are highly similar to their *in vivo* counterparts (Rossi et al., 2020), which is likely due to their recapitulation of developmental trajectories and their spatiotemporal response to mechano-chemical signals from proximal tissues. In this work, we showed that gastruloids expressed key hematopoietic genes in a temporally relevant fashion. The gastruloids also spontaneously expressed *Hoxb4* in the hematoendothelial and endothelial clusters. Gastruloids also implement collinear *Hox* transcriptional patterns along the anteroposterior axis (Beccari et al., 2018) and form morphologically relevant organ primordia, in a similar fashion to the tissue spatial organization in embryos (Rossi et al., 2020). Furthermore, numerous *in vitro* and *in vivo* experiments have previously revealed that, within the hemogenic endothelium, a Flk1^+^ precursor population has the ability to undergo endothelial-to-hematopoietic transition (Jaffredo et al., 1998; Ottersbach, 2019; Zovein et al., 2008). Such *Flk1*^+^ precursors were also detected in gastruloids, both by scRNA-seq and flow cytometry analysis, together with the expression of other canonical hematoendothelial markers.

We have previously shown that gastruloids develop an endodermal primitive gut tube-like structure (Vianello and Lutolf, 2020), which forms close to the cardiac compartment (Rossi et al., 2020). It is known that endoderm-derived factors are needed to guide hemogenic fate and HSC development (Baron, 2003; Belaoussoff et al., 1998; Miura and Wilt, 1969; Pardanaud and Dieterlen-Lièvre, 1999; Peeters et al., 2009; Wilt, 1965). We hypothesized that factors secreted by the endodermal compartment in gastruloids influence the development and localization of hemogenic cells. Indeed, we have described the emergence of CD41^+^ blood progenitor cells that arise in the anterior portion of gastruloids, in close proximity to the vascular-like network and the endodermal compartment. Furthermore, these evolving hematopoietic progenitor cells display multipotent clonogenic potential in CFU assays *in vitro* and *in vivo* at later stages of gastruloid development. Importantly, we observed the emergence of erythroid cells that are the first differentiated hematopoietic cells to appear in gastruloids similar to the embryonic development (Costa et al., 2012). We have confirmed this by the expression of embryonic hemoglobin isoforms in gastruloids at 168 h. Intriguingly, we observe the expression of these genes within the endothelial cluster, and mainly in a sub-cluster that we previously associated with endocardial development (Rossi et al., 2020). This suggests that these cells might arise from the hemogenic endocardium, which is a known site for hematopoiesis at the early heart tube developmental stage (Nakano et al., 2013).

In addition to genetic and morphological studies, we compared different cell lines through flow cytometry analysis to corroborate our findings. We observed a similar trend in the expression of hematopoietic surface markers across distinct cell lines at different time points. However, we observed differences in the propensity of each cell line to generate cells that expressed typical blood progenitor markers. Our observations are in line with previous studies that suggest that the genetic background of the used cell line can skew mESC differentiation (Ortmann et al., 2020) and gastruloid cell-type diversity (van den Brink et al., 2020).

In conclusion, our data suggest that gastruloids are excellent candidates to model hematopoietic development *in vitro* in an embryo-like context. Gastruloids can serve as model systems to study the emergence of blood progenitors from different embryonic locations, to recapitulate key signals and developmental events, all with spatiotemporal fidelity.

In the future, gastruloids can be used to model the emergence of hematopoietic precursor cells with the potential to decipher their cellular origins and complex interactions of blood precursors with the proximal endothelium. The simple scalability and facile manipulability of gastruloids provides the potential for large-scale production of hematopoietic progenitors in an embryo-like context. Since human PSC-derived gastruloids have recently been described (Moris et al., 2020), future studies are likely to transfer this know-how to human models and open new avenues for human *in vitro* developmental studies, screening, and personalized medicine approaches. Indeed, previous attempts to generate HSCs in culture were mainly based on the use of embryoid bodies (EBs), through ectopic expression of key hematopoietic genes (Kyba et al., 2002; Wang et al., 2005), or a combination of morphogen-driven directed differentiation and transcription factor-mediated cell fate conversion (Lu et al., 2016; Sugimura et al., 2017). Although these protocols have shown that transplantable cells can be generated, they are not ideal for capturing early developmental events with spatiotemporal fidelity. Moreover, ectopic expression of key hematopoietic genes represents a developmental shortcut that could bypass crucial steps in HSC development. Our developmental biology-inspired approach may instead lead to the generation of cells and progenitors with improved therapeutic relevance in the future. Finally, we envision that gastruloids have applicability beyond studies of cardiovascular and hematopoietic development and serve as *in vitro* models for other mechanistic studies of embryonic developmental biology.

## Material and Methods

### Cell Culture

mESCs were cultured as previously described (Rossi et al., 2020). Briefly, all cells were cultured at 37°C, 5% CO2 in growth medium (DMEM, 10% Embryonic Stem Cell qualified FBS (Gibco), NEAA, Sodium Pyruvate, β-mercaptoethanol, 3μM CHIR99021 (Chi), 1μM PD025901 and 0.1μg ml-1 LIF). *Gata6-Venus* (Freyer et al., 2015) and *Flk1-GF*P (Jakobsson et al., 2010) cells were cultured on gelatinized tissue-culture flasks; *Sox1-GFP::Brachyury-mCherry* cells (Deluz et al., 2016) without any coating. All cells were routinely tested for Mycoplasma with Mycoalert plus mycoplasma detection kit (Lonza) or by PCR.

### Gastruloid culture

Gastruloids were generated as previously described (Baillie-Johnson et al., 2015; Rossi et al., 2020; van den Brink et al., 2014). Briefly, 300-700 mESCs were aggregated in 40μl N2B27 in 96-well Clear Round Bottom Ultra-Low Attachment Microplates (7007, Corning). After 48h, 150μl per well of 3μM Chi in N2B27 were added. At 72h, 150μl of medium were removed and substituted with 150μl of fresh N2B27. From 96h, the medium was changed to N2B27+++, which contains 30ng ml-1 bFGF (PMG0034, Gibco), 5ng ml^−1^ VEGF 165 (PHC9394, Gibco) and 0.5mM L-ascorbic acid phosphate (013-12061, Wako). From 120h on, half of the medium was changed daily. From 144h, N2B27 was used for daily medium changes. For immunofluorescence analysis, gastruloids at 96h were transferred in Ultra-Low Attachment 24-Well Plates (3473, Corning) with 100μl of medium, plus 700μl of fresh N2B27+++, and cultured on an orbital shaker placed at 37°C, 5% CO2 at 100rpm (VWR mini shaker), with the same culture schedule.

### Whole mount immunofluorescence and light-sheet imaging

Immunofluorescence on whole-mount gastruloids was performed as previously described (Baillie-Johnson et al., 2015; Rossi et al., 2020). Briefly, gastruloids were washed with PBS and fixed in 4% PFA for 2h at 4°C on a rotating shaker. Samples were repeatedly washed with PBS and blocking buffer (PBS, 10%FBS, 0,2%Triton X-100), then blocked for 1h at 4°C with blocking buffer. Gastruloids were then incubated O/N with primary antibodies in blocking buffer, at 4°C on a rotating shaker. On the following day, gastruloids were repeatedly washed with blocking buffer, at 4°C while shaking, and incubated O/N with secondary antibodies and DAPI (2μg ml-1, Sigma-Aldrich) in PBS, at 4°C on a rotating shaker. On the following day, gastruloids were washed for 1h with blocking buffer, at 4°C on a rotating shaker, and rinsed in PBS, 0,2% FBS, 0,2% Triton X-100. For light-sheet imaging, gastruloids were mounted in 1% low melt agarose in glass capillaries, and cleared with CUBIC-mount solution (Lee et al., 2016) O/N. Imaging was performed in CUBIC-mount solution with a Zeiss Light-sheet Z1 microscope equipped with a Plan-Neofluar 20x/1.0 Corr nd=1.45 objective. Images were processed using Imaris software for 3D image analysis.

The following primary antibodies were used: rabbit anti-CD41 (ThermoFischer Scientific, PA5-79526, 1:200); rabbit anti- CD34 (clone EP373Y, Abcam, 1:100); goat anti-Sox17 (R&D, AF1924, 1:200); rat anti CD31 (BD, MEC13.3, 1:100). The following secondary antibodies were used: donkey anti rabbit 647 (ThermoFischer Scientific, 1:500); goat anti-rat 568 (ThermoFischer Scientific, 1:500), donkey anti rabbit 568 (ThermoFischer Scientific, 1:500), donkey anti rat 647 (ThermoFischer Scientific, 1:500). DAPI (2μg ml-1, Sigma-Aldrich) was used to stain nuclei.

### Immunofluorescence on sections

For immunofluorescence on tissue sections, gastruloids were washed in PBS and fixed O/N in 4% PFA, at 4°C on an orbital shaker. The following day, samples were washed in PBS and embedded in HistoGel (ThermoFisher). HistoGel blocks were then processed with a Tissue-Tek VIP 6 AI Vacuum Infiltration Processor (Sakura) and embedded in paraffin. 4μm paraffin sections were obtained with a Hyrax M25 microtome (Zeiss). Slides were processed through de-waxing and antigen retrieval in citrate buffer at pH6.0 (using an heat-induced epitope retrieval PT module, ThermoFischer Scientific) before proceeding with immunostaining. Sections were then blocked and permeabilized for 30min in 1% BSA, 0.2% Triton X-100 in PBS and blocked for 30min in 10% goat or donkey serum (Gibco) in PBS at RT. Primary antibodies were incubated O/N at 4°C in PBS, 1.5% donkey or goat serum. On the following day, slices were washed twice in 1% BSA, 0.2% Triton X-100 in PBS and incubated with secondary antibodies at RT for 45min. Finally, slices were washed twice in 0.2% Triton X-100 in PBS and mounted with Fluoromount-G (SouthernBiotech). Pictures were acquired with an upright Leica DM5500 scanning microscope equipped with a CCD DFC 3000 black and white camera. The following primary and secondary antibodies were used: rabbit anti-CD41 (ThermoFischer Scientific, PA5-79526, 1:200); donkey anti rabbit 647 (ThermoFischer Scientific, 1:500). DAPI (2μg ml-1, Sigma-Aldrich) was used to stain nuclei.

### Analysis of single cell-RNAseq data

Single cell analysis was performed with Seurat v3.1 on the dataset described in (Rossi et al., 2020) and available at NCBI GEO (http://www.ncbi.nlm.nih.gov/geo/) under the accession number GSE158999, using codes described in (Rossi et al., 2020) and available at (https://github.com/nbroguiere/Cardiac_Gastruloids), focusing on a close-up view of clusters relevant for this paper (as shown in **Figure 1**).

### RNA extraction and qRT-PCR

RNA extraction was performed using the RNeasy Micro kit (Qiagen), according to manufacturer’s instructions. RNA was quantified with a Nanodrop (ND-1000). 1μg of RNA was reverse-transcribed using the iScript cDNA Supermix kit (Biorad). 1.5μl of 1:5 diluted cDNA was used per reaction, in a total volume of 10μl. 384 plates were loaded with an automated liquid handling system (Hamilton Microlab Star). qPCR was performed with a 7900HT Fast PCR machine (Applied Biosystems), using Power SYBR Green PCR Master Mix (Applied Biosystems), with an annealing temperature of 60°C. β-actin expression was used to normalize data. Relative fold expression was calculated with the 2−ΔΔCT method. 500nM of the following primers were used: β-actin FOR CTGTCGAGTCGCGTCCACC; β-actin REV CGCAGCGATATCGTCATCCA; Gata1 FOR GTGGCTGAATCCTCTGCATCA; Gata1 REV TAAGGTGAGCCCCCAGGAAT; Gata2 FOR GCCGGGAGTGTGTCAACTG; Gata2 REV AGGTGGTGGTTGTCGTCTGA; Mecom FOR AATAAATCCGAAACGCGTGGT; Mecom REV CCTACATCTGGTTGACTGGCA; Runx1 FOR AGGCAGGACGAATCACACTG; Runx1 REV CTCGTGCTGGCATCTCTCAT; Tal1 FOR CACTAGGCAGTGGGTTCTTTG; Tal1 REV GGTGTGAGGACCATCAGAAATCT; Hoxb4 FOR AAAGAGCCCGTCGTCTACC; Hoxb4 REV AGTGTAGGCGGTCCGAGAG; Hbb-bh1 FOR GGAAACCCCCGGATTAGAGC; Hbb-bh1 REV CTGGGGTGAATTCCTTGGCA; Hbb-b1 FOR TGTCTCTTGCCTGTGGGGAA; Hbb-b1 REV GAAATCCTTGCCCAGGTGGT; Hbb-b2 FOR TGAAGGCCCATGGCAAAAAG; Hbb-b2 REV CGATCGCATTGCCTAGGAGC; Hba-a1 FOR CTGAAGCCCTGGAAAGGATGT; Hba-a1 REV AGAGCCGTGGCTTACATCAAA; Hba-x FOR ATCAGGCCAGTCTTGAGTGC; Hba-x REV GGAGCTTGAAGTTGACCGGA; Hbb-y FOR TCTGCCATAATGGGCAACCC; Hbb-y REV TGCCGAAGTGACTAGCCAAA.

### FACS analysis and cell sorting

For FACs analysis and cell sorting, gastruloids were washed once in PBS and digested in 4mg ml^−1^ dispase I (Roche), 3mg ml^−1^ collagenase IV (Gibco) and 100μg ml^−1^ DNase I (Roche) in PBS (2 digestion cycles at 37°C, 4 min each, each followed by mechanical dissociation through gentle pipetting). Digestion was blocked with 10% FBS. Samples were then centrifuged, resuspended in sorting buffer (PBS, 10% FBS, 1mM EDTA, 1% P/S) and incubated for 45 min with antibodies and 10 min with DAPI (2μg ml^−1^, Sigma-Aldrich), always on ice. Unstained, fluorescent minus one (FMO) and single-color samples were used as controls. Samples were analyzed with a BD LSR Fortessa flow cytometer. Cell sorting was performed with the Aria Fusion cell sorter (BD Bioscience). The following antibodies and dilutions were used: CD34 eFluor660 Mouse Rat IgG2a kappa (eBioscience 50-0341-82, cloneRAM34) 1:80; Sca1 BV711 Mouse Rat IgG2a kappa (Biolegend 108131, clone D7) 1:80; cKit PE-Cy5 Mouse Rat IgG2b kappa (eBioscience 15-1171-82, clone2B8) 1:640; CD31 PE Mouse Rat IgG2a kappa (BD Pharmingen 561073, clone MEC13.3) 1:1280; CD48 BV421 Mouse Amerian Hamster IgG (Biolegend 103427, clone HM48-1) 1:640; CD41 BV605 Mouse Rat IgG1 kappa (BD Pharmingen 747728, clone MWReg30) 1:80; TER119 APC-Cy7 Mouse Rat IgG2b kappa (Biolegend 116223, clone TER-119) 1:160; CD93 PE-Cy7 Mouse Rat IgG2b kappa (Biolegend 136506, clone AA4.1) 1:640; CD45 AF700 Mouse Rat IgG2b kappa (Biolegend 103128, clone 30-F11) 1:30.

### Colony-forming unit assay

To test their colony formation capacity (CFC), cells with presumptive hemogenic capacity were sorted from day 7 gastruloids based on surface marker expression. Sorted cells were added at serial dilutions to 1.5ml of MethoCult (Stem Cell Technologies, H4434). The mixture of MethoCult and cells was vortexed and transferred with a blunt end needle syringe (Stem Cell Technologies, 28110) into meniscus-free SmartDish plates (6-well-plate, Stem Cell Technologies, 27370). To allow constant humidification, the 6-well plates were placed in a large 150mm square dish (Corning, CLS431111) containing small 3.5×10 mm tissue culture dishes filled with water. The cultures were incubated at 37°C, 5% CO2 for 14 days and the CFC content was automatedly imaged after 7 and 14 days by using the StemVision imaging system (Stem Cell Technologies, Version 2.0.1.0). Additionally, higher-resolution images of colonies were acquired with a PALM Microbeam microscope (Zeiss) equipped with an AxioCamICc1 color camera and 5x and 10x objectives.

### Transplantation and Spleen assay

For transplantation, *Gata6-Venus* (CD45.2) Gastruloids at 168h were dissociated as described for FACs analysis. 10 weeks old, female, Bl6, CD45.1 recipient mice were lethally irradiated 24 hours before transplantation with a split dose of 2x 425cGy. 5×10^6^ dissociated gastruloid cells were injected in the mice tail vein together with 2×10^4^ competitor bone marrow cells isolated from Bl6, CD45.1/2 mice, in a total volume of 200 µl. Control mice received just 2×10^4^ competitor bone marrow cells isolated from Bl6, CD45.1/2 mice, in a total volume of 200 µl. Mice were euthanized 11 days after transplantation, and spleens were collected for analysis. Whole spleen images were acquired with a Leica MZ16 1FA stereomicroscope equipped with Leica CLS 150X illumination and DFC480 color camera. All mice were purchased from Charles River Laboratories International and maintained at EPFL conventional animal facility, in microisolator cages, provided with food and water *ad libitum*. Animal experiments were performed in compliance with the Swiss law after approval from the local and federal authorities (license VD2242.2).

### Statistics

All data in graphs are represented as mean ± SD. Statistical analysis between two columns was performed using two-tailed unpaired Student’s t test. Statistical significance was calculated using the Graphpad Prism software, that was also used to generate all graphs.

*P < 0.05; **P < 0.01; ***P < 0.001; confidence intervals 95%; alpha level 0.05.

## Acknowledgements

We thank all members of the Lutolf laboratory for helpful discussions, especially Nicolas Broguiere for his help with single-cell RNAseq. We thank Alfonso Martinez Arias (UPF Barcelona), Christian Schröter (MPI Dortmund), Alexander Medvinsky (MRC Edinburgh) and David Suter (EPFL) for providing reporter cells. We thank all members of EPFL Bioimaging and Optics Facility, Histology Core Facility, Flow Cytometry Core Facility, and Gene Expression Core Facility for their technical support. This work was funded by a Sinergia grant (CRSII5_189956) from the Swiss National Science Foundation, the National Center of Competence in Research (NCCR) Bio-Inspired Materials, and EPFL.

---

**Supplementary Figure 1.**
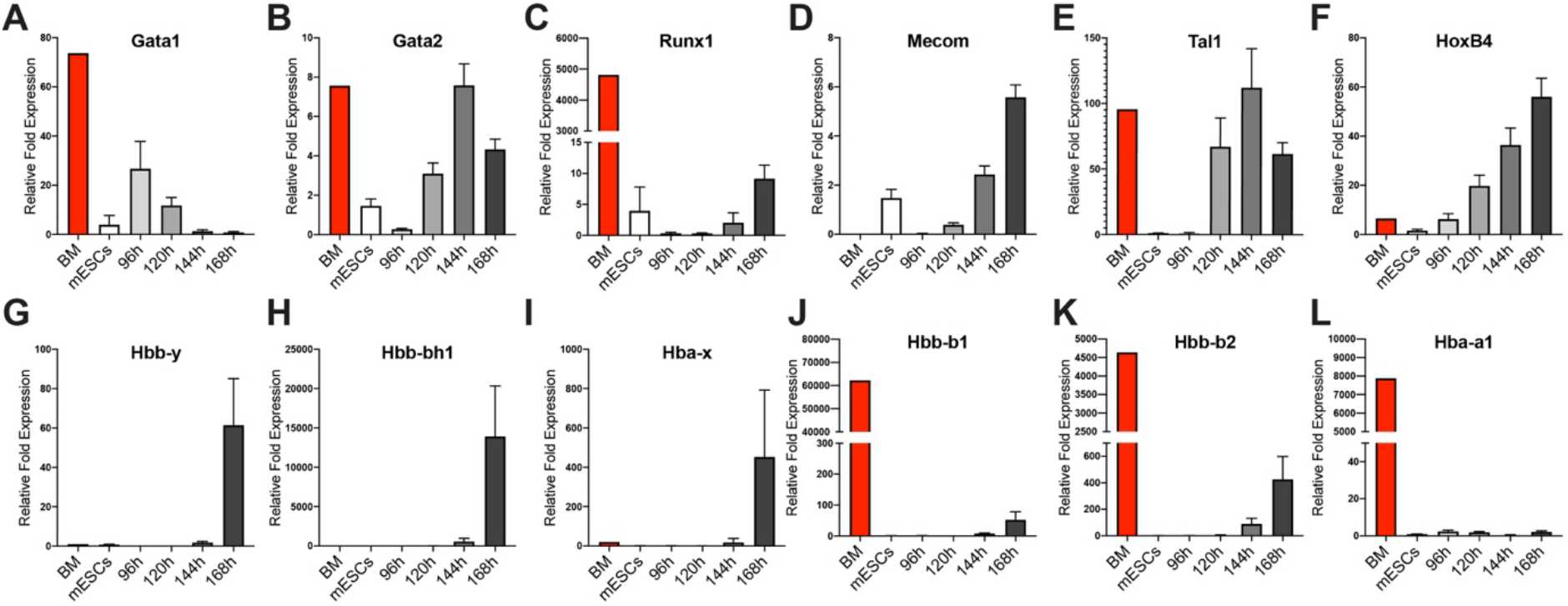
Gastruloids express markers of early blood development. **A-F**, qRT-PCR showing the relative fold expression of genes associated with hematopoietic development in *Sox1- GFP::Brachyury-mCherry* gastruloids from 96h to 168h, compared to mESCs. **G-L**, qRT-PCR showing the relative fold expression of embryonic (**G-I**) and adult (**J-L**) hemoglobin genes in *Sox1- GFP::Brachyury-mCherry* gastruloids from 96h to 168h, compared to mESCs. BM, bone marrow. Data are expressed as ± SD, n=4 replicates. *Related to Figure 1.*

**Supplementary Figure 2.**
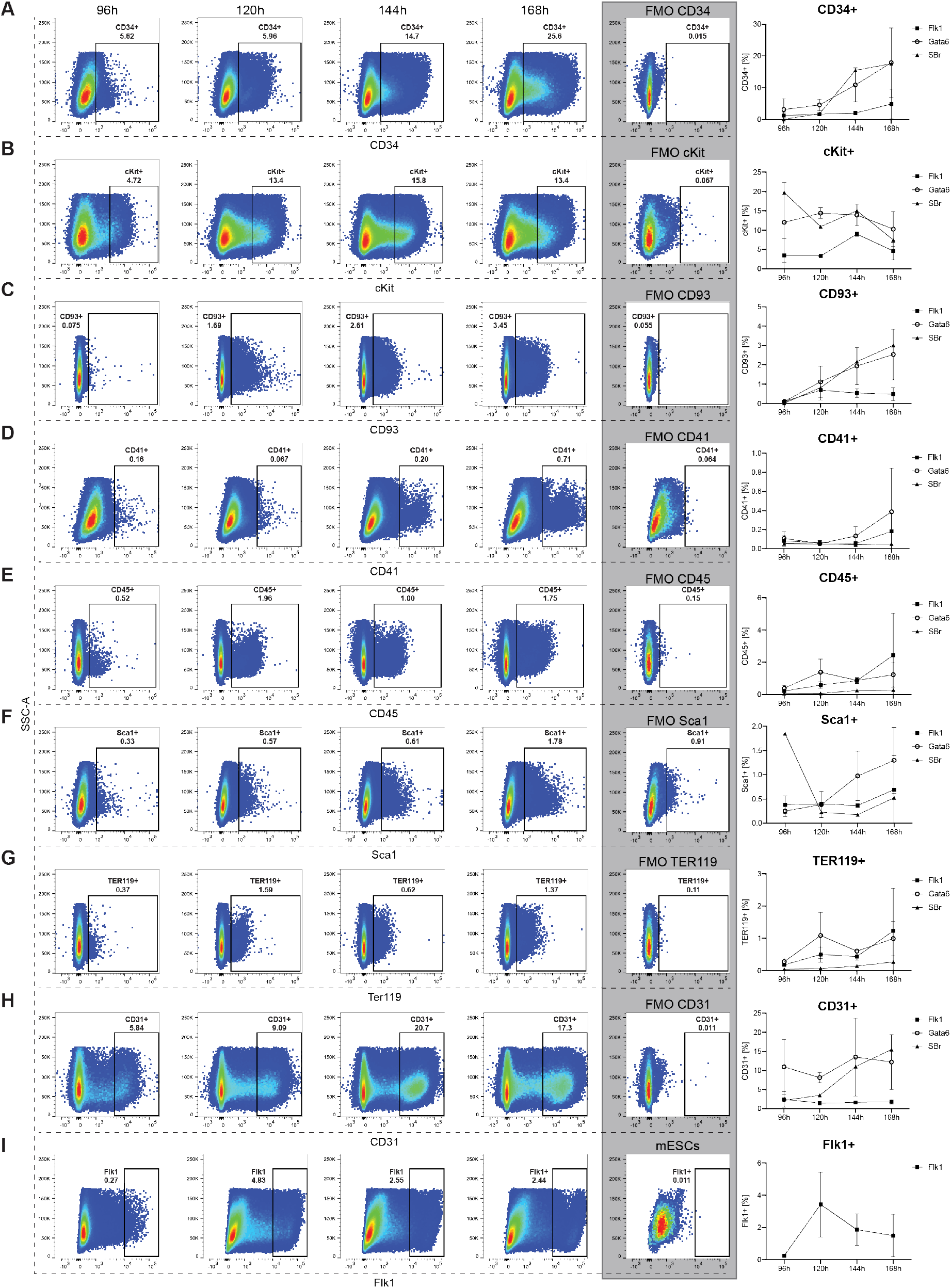
Characterization of hematopoietic surface markers in gastruloids. **A-I**, Representative FACS plots and relative quantification showing CD34 (A), cKit (B), CD93 (C), CD41 (D), CD45 (E), Sca1 (F), Ter119 (G), CD31 (H), and Flk1 (I) expression in gastruloids from 96h to 168h as well as the relative FMO controls (shaded in grey). *Flk1-GFP* (n = 2 replicates), *Gata6-Venus* (n = 2 replicates) and *Sox1-GFP::Brachyury-mCherry* (n = 1 replicate). Data are represented as mean ± SD. *Related to Figure 2.*

**Supplementary Figure 3.**
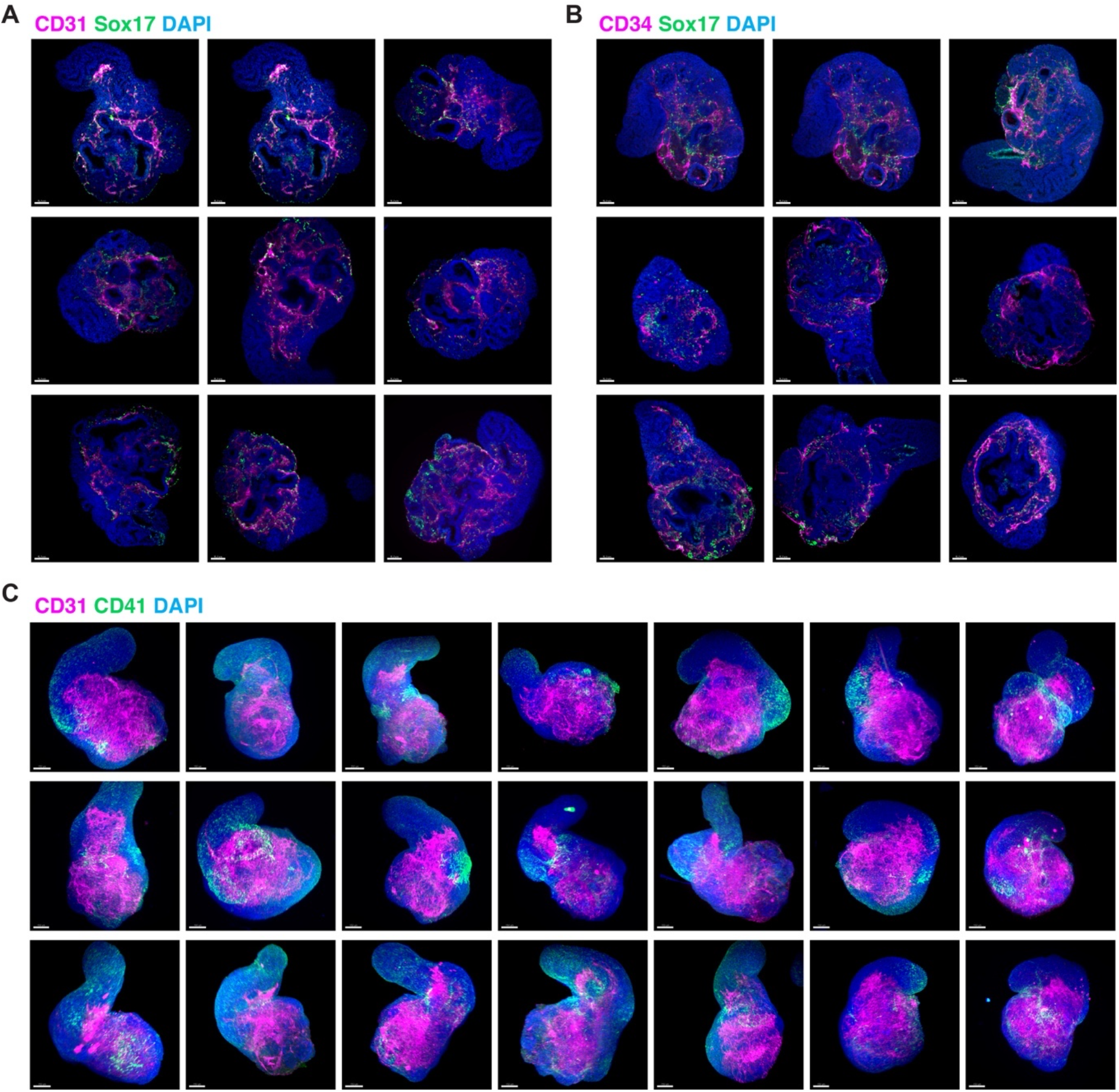
Emergence of blood progenitors in gastruloids. **A**, Representative collection of light-sheet images of *Gata6-Venus* gastruloids at 168 h showing the co-expression of CD31 and Sox17 in single Z-planes. **B**, Representative collection of light-sheet images of *Gata6-Venus* gastruloids at 168 h showing the co-expression of CD34 and Sox17 in single Z-planes. **C,** Representative collection of light-sheet images of *Gata6-Venus* gastruloids at 168 h showing CD41^+^ clusters arising in close proximity to the CD31^+^ vascular network. Scale bars, 100μm. *Related to Figure 3.*

**Supplementary Figure 4.**
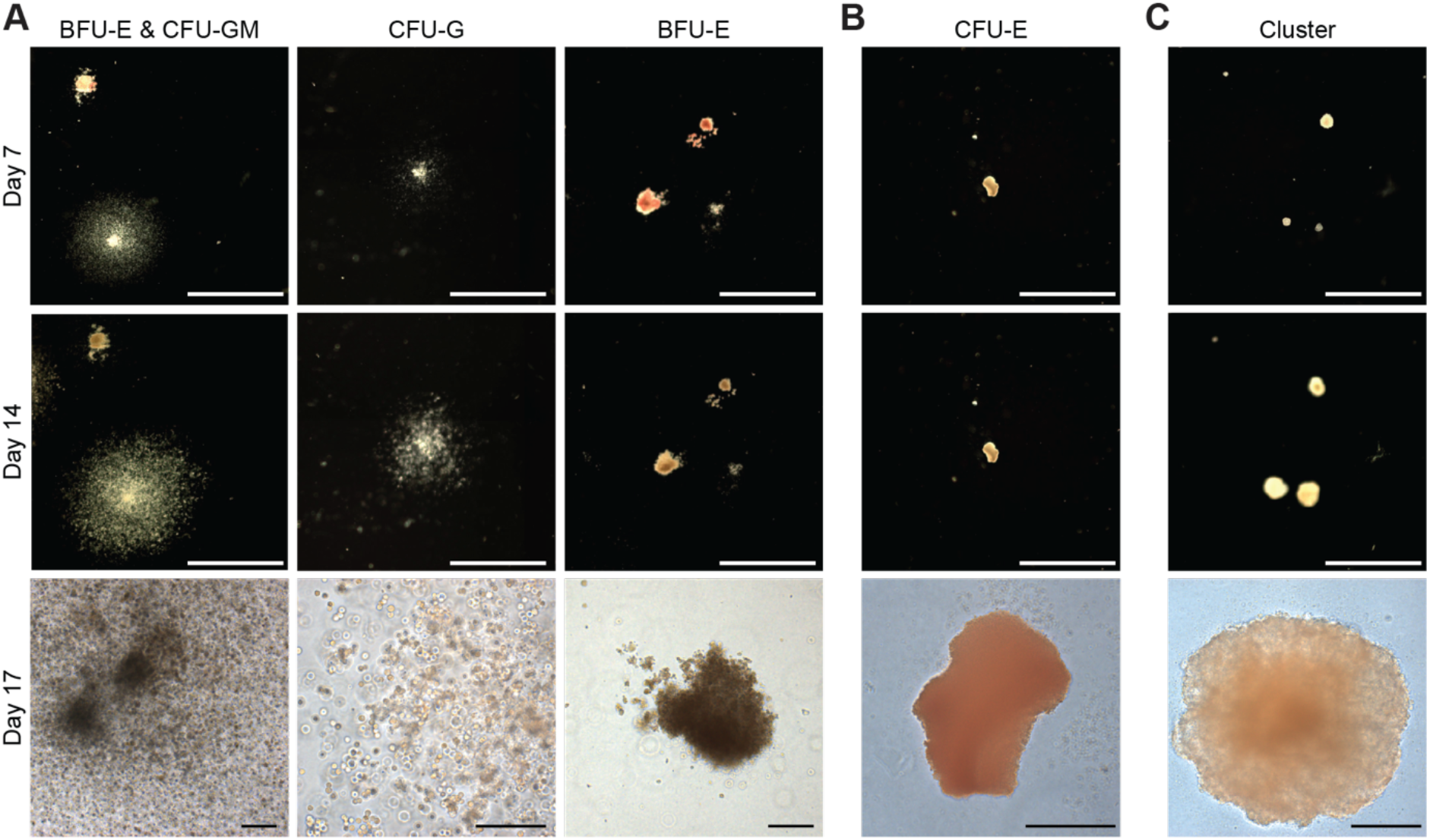
Multilineage clonogenic potential of gastruloid-derived blood progenitors. **A**, Representative images of BFU-E, CFU-GM and CFU-G colonies derived from cKit^+^/CD34^+^/TER119^−^/CD41^+^ cells. **B**, Representative images of CFU-E colonies derived from the Not(cKit^+^/CD34^+^)/TER119^+^/CD41^−^ population. **C**, Representative images of cluster formation of Gata6-Venus mESCs. All cell populations were cultured in Methocult and imaged after 7, 14, and 17 days. BFU-E, burst-forming-unit-erythroid; CFU-GM, colony forming unit-granulocyte, macrophage; CFU-E, colony forming unit-erythroid. Scale bars 200μm. *Related to Figure 4.*

